# Single-Molecule Analysis of DNA-binding proteins from Nuclear Extracts (SMADNE)

**DOI:** 10.1101/2022.04.01.486290

**Authors:** Matthew A. Schaich, Brittani L. Schnable, Namrata Kumar, Vera Roginskaya, Rachel C. Jakielski, Roman Urban, Zhou Zhong, Neil M. Kad, Bennett Van Houten

## Abstract

Single-molecule characterization of protein-DNA dynamics provides unprecedented mechanistic details about numerous nuclear processes. Here, we describe a new method that rapidly generates single-molecule information for fluorescently tagged proteins isolated from nuclear extracts of human cells. This approach determines binding lifetimes (*k*_off_), events per second (∼*k*_on_), positional dependence (specificity), and characterizes 1D diffusion along DNA. We demonstrated the wide applicability of this approach on three forms of DNA damage using seven native DNA repair proteins and two structural variants, including: poly(ADP-ribose) polymerase (PARP1), heterodimeric ultraviolet-damaged DNA-binding protein (UV-DDB), xeroderma pigmentosum complementation group C protein (XPC), and 8-oxoguanine glycosylase 1 (OGG1). By measuring multiple fluorescent colors simultaneously, we additionally characterized the assembly and disassembly kinetics of multi-protein complexes on DNA. Thus, Single-Molecule Analysis of DNA-binding proteins from Nuclear Extracts (SMADNE) provides new insights about damage recognition and represents a universal technique that can be used to rapidly characterize numerous protein-DNA interactions.

## Introduction

Watching DNA-binding proteins interact with DNA substrates in real-time at the single-molecule level illuminates how proteins detect and bind their targets at extraordinary detail. Key information about binding stoichiometry, order of assembly and disassembly, and how proteins diffuse to find their DNA targets are gained through single molecule analysis^1-3^. Various imaging techniques and optical platforms have been employed to resolve fluorescent proteins to the single-molecule level, but most of these techniques cluster into two broad categories – studies performed with purified proteins with defined conditions ^4-6^ or studies performed in living cells ^7-9^.

In single-molecule fluorescence studies of DNA-binding proteins, the molecules of interest must first be purified and then be labeled with a fluorescent tag, ranging in size from small chemical dyes to fluorescent proteins to large quantum dots (Qdots) ^7^. These techniques hold the distinct advantage of knowing precisely what proteins are binding to the DNA substrates of interest held in a static location. However, overexpressing, purifying, and labeling some proteins can prove difficult due to loss of activity. In addition, even using Qdots conjugation with antibodies, labeling is less than 100%^10^. Furthermore, other protein factors that may contribute to stabilizing or destabilizing ligand binding and/or catalytic activity are lost during purification. The resulting studies of purified DNA-binding proteins may therefore not accurately represent how these proteins work in the context of the complex cellular milieu of the nucleus.

Conversely, single-molecule studies of DNA-binding proteins have also been performed within living cells ^7-9^. These techniques developed for prokaryotes initially, but recent work has allowed for this imaging even in mammalian cells ^11-16^. While these approaches are the most biologically relevant, watching DNA-binding proteins sort through the complex genome to find their specific binding sites has proven challenging, but technically possible ^17^. However, these approaches rely on having low enough fluorescence signal to resolve individual proteins, and therefore there are often many unlabeled proteins of interest competing and altering binding lifetimes. Furthermore, proteins diffusion along DNA cannot be studied when DNA strand orientation is unknown.

To overcome many of the challenges associated with the single-molecule characterization of DNA-binding proteins, we designed a new method that exists at the confluence of these two types of techniques, which we have termed SMADNE for <u>S</u>ingle-<u>M</u>olecule <u>A</u>nalysis of <u>D</u>NA-binding Proteins from <u>N</u>uclear <u>E</u>xtracts. SMADNE applies similar principles of previous single-molecule work with cellular extracts ^18-25^ while making several significant improvements, allowing application to human cells and scalability to numerous proteins that bind DNA. Using the LUMICKS C-trap combined optical tweezers, microfluidics, and 3-color confocal microscope, we precisely define the positions of fluorescently-tagged DNA repair proteins on 48.5 kb DNA substates containing defined types of damage. As shown below, SMADNE provides binding specificity and diffusivity measurements including characterizing multiple proteins simultaneously binding DNA damage with over 4 orders of magnitude of duration (0.1 to >100 s) and a wide range of 1D diffusivity values (from 0.001 to 1 μm^2^ s^-1^), with similar precision as other single molecule techniques^4,26-34^. At the same time, SMADNE bridges the complex milieu of the nuclear environment containing thousands of proteins to a system where fluorescently tagged single particles can be followed and characterized. Thus, SMADNE has broad applicability to provide detail mechanistic information about diverse protein-DNA and protein-protein interactions.

## Results

### SMADNE workflow and characterization of PARP1 binding to damaged DNA

To study fluorescently tagged DNA-binding proteins from nuclear extracts, we developed the workflow shown in Fig. 1a and b. Using the LUMICKS C-trap optical traps, streptavidin coated polystyrene beads were captured and biotinylated 48.5 kb DNA was suspended between the beads (Fig 1c, left panel). After flowing in the nuclear extract containing the fluorescently labeled protein of interest, flow was stopped, and 2D confocal images were collected to verify binding of the protein to the DNA (Fig. 1c, middle panel). Then, the area being scanned was reduced to only the central DNA position. In this 1-dimensional scanning mode, imaging rates as fast as 6 msec per scan can be achieved. These data appear as fluorescent time streaks (kymographs) showing the fluorescently-tagged protein position over time, where the Y-axis represents the position on the DNA and the X-axis shows the scan time (Fig. 1c, right panel). In this mode, the Y-axis represents positions on the DNA where binding occurs, and the X-axis shows the scan time, which in this kymograph is 30 msec increments.

**Fig. 1:**
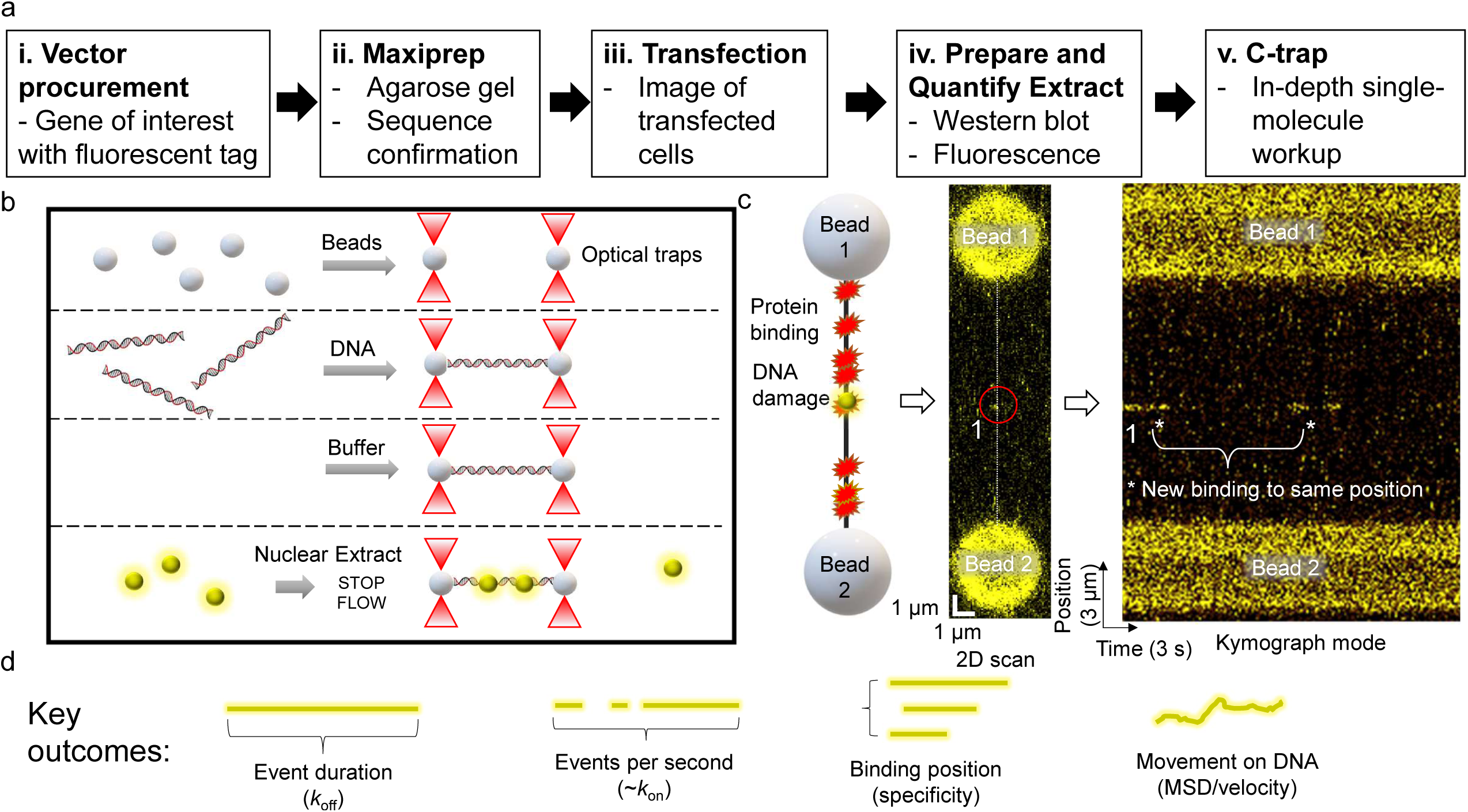
The workflow and experimental outcomes of SMADNE. **a**, SMADNE workflow. **b**, A diagram of the imaging techniques using four channels separated by laminar flow. **c**, A cartoon of a DNA substrate for SMADNE suspended between two polystyrene beads and tagged proteins (yellow spheres) binding to sites of DNA damage. This substrate (nicked DNA) is also shown as a 2D scan (one YFP-PARP1 binding event numbered and circled) and in kymograph mode (numbered spot marked). Event one dissociated before the kymograph started and then another event appeared at the same position later (asterisks). Binding events appear as lines in the kymograph because time is indicated on the X axis and position on the Y axis. **d**, The four major outcomes obtained from SMADNE characterization.

To validate the general utility of SMADNE, we examined series of fluorescently tagged-DNA repair proteins on various DNA substrates, namely poly(ADP-ribose) polymerase 1 (PARP1), xeroderma pigmentosum complementation group C protein (XPC), apurinic/apyrimidinic endonuclease 1 (APE1), DNA polymerase β (Pol β), DNA damage-binding protein 1 (DDB1) and DNA damage-binding protein 2 (DDB2). In Fig. 1, YFP-PARP1 forms transient complexes on DNA nicked with a site-specific nickase creating time streaks in the kymograph mode. Notice how multiple molecules revisit the same positions on the DNA (Fig. 1c, asterisks). These likely represent multiple events on the same damage site. Using SMADNE, we determined four key outcomes: 1) how long a binding event lasts from start to finish (*k*_off_); 2) how many binding events per second occur (related to *k*_on_); 3) the position of binding events along the DNA; and 4) how bound proteins diffuse along the DNA (Fig. 1d). For YFP-PARP1 at 10 pN of DNA tension, the average lifetime exhibited was 4.3 seconds, events occurred at 0.13 events per second, the positions agreed with the expected sites, and no diffusion along the DNA was observed (Fig. 2).

**Fig. 2:**
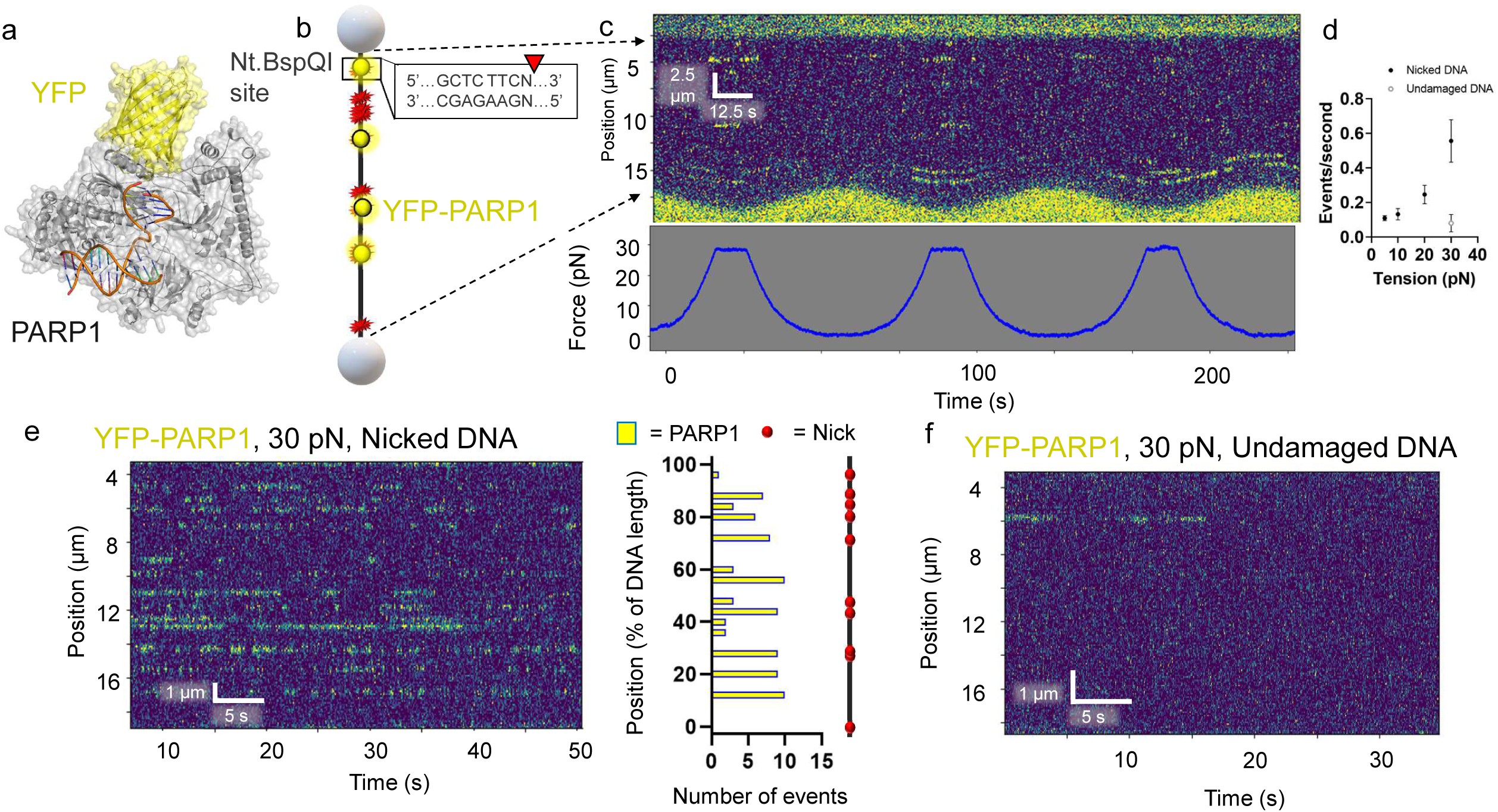
DNA tension influences DNA nick detection by PARP1. **a**, A structural model of PARP1 bound to nicked DNA with YFP tag (PDB codes 3ED8 and 4KLO) generated as in this reference^1^. **b**, A schematic of the DNA suspended between streptavidin beads containing 10 discrete nicks from the nickase Nt.BspQI. **c**, An example kymograph of PARP1 binding DNA at oscillating tensions from 5 pN to 30 pN. Binding events shown in yellow and tension measurements shown below in blue. **d**, Number of events per second at various DNA tensions held constant. Error bars represent the SEM of three experiments. Gray circle represents undamaged DNA. **e**, An example kymograph of PARP1 binding DNA at constant tension (30 pN). Positional analysis shown to the right showed biding at the expected sites, but also several sites that were bound multiple times that did not contain the recognition sequence by Nt.BspQI. **f**, Undamaged DNA exhibited reduced YFP-PARP1 binding, even at 30 pN.

### SMADNE characterization of PARP1 binding nicked DNA: increasing DNA tension increased binding events

To demonstrate the broad applicability of SMADNE to various DNA repair proteins and different forms of DNA damage, the binding interactions were examined for YFP-tagged PARP1 from nuclear extracts on DNA containing ten nicks generated by a sequence-specific nickase (Fig. 2a and b). Unexpectedly, increasing the tension on the DNA from 5 pN to 30 pN dramatically increased the number of YFP-PARP1 events per second. At 30 pN, new binding sites also appeared that were not observed at lower tension (Fig. 2c). It is possible that the higher tension makes previously existing nicks more identifiable by PARP1. Datasets were then collected at various constant DNA tensions. While binding lifetimes stayed relatively consistent as analyzed by fitting a cumulative residence time distribution (CRTD) to an exponential decay function, events per second increased 4-fold at 30 pN of tension. In contrast, undamaged events per second remained low even at high tensions (Fig. 2d). YFP-PARP1 from nuclear extracts repeatedly bound at specific locations on the DNA, both on undamaged and damaged DNA (Fig. 2e, f) indicating repeated specific binding events occurred at the nick sites. Datasets collected at 30 pN tension resulted in numerous binding events at 13 positions on the nicked DNA, indicating some off-target DNA damage present in our DNA sequence (Fig. 2e). While no previous work to our knowledge has examined PARP1 binding to nicked DNA at the single molecule level, we previously examined single molecules of purified PARP1 labeled with Qdot binding to abasic sites, finding that PARP1 largely bound its substrate via 3D diffusion, which agrees with the results we observed with nicked DNA using SMADNE^4^.

### Application of SMADNE to study transient DNA interactions

We next sought to push the limits of the SMADNE technique to more transient interactions, such as XPC-RAD23B that diffuses along the DNA while detecting UV damage, as well as APE1 or Pol β binding to nicks with much lower affinity (Fig. 3) ^35-37^. For XPC-RAD23B, eGFP-tagged XPC and untagged RAD23B were co-transfected, and eGFP signal was observed on UV-damaged (40 J/m^2^) DNA (Fig. 3a). Thus, XPC binds UV-damaged DNA and, in 44% of events, diffused along the DNA (Fig. 3b, d). Binding lifetimes for XPC in nuclear extracts were similar to those observed for purified XPC^36^, with the CRTD fitting to a double exponential to yield one lifetime at 48.6 seconds and a second lifetime at 0.89 seconds, with the fast component contributing 67 percent (Fig. 3c). Mean squared dissociation (MSD) analysis performed on the motile XPC molecules (Fig. 3e) revealed a diffusion constant with a geometric mean of ∼ 0.03 µm^2^ s^-1^, which agrees with previously published work (Fig. 3f). Additionally, tGFP-tagged APE1 and Pol β binding were also characterized on DNA with 10 nicks as previously done with PARP1. Both proteins bound the nicked substrate with relatively lower affinity, with APE1 exhibiting a binding lifetime of 0.3 s (Fig. 3g-i) and Pol β binding for 1.8 s (Fig. 3j-l).

**Fig. 3:**
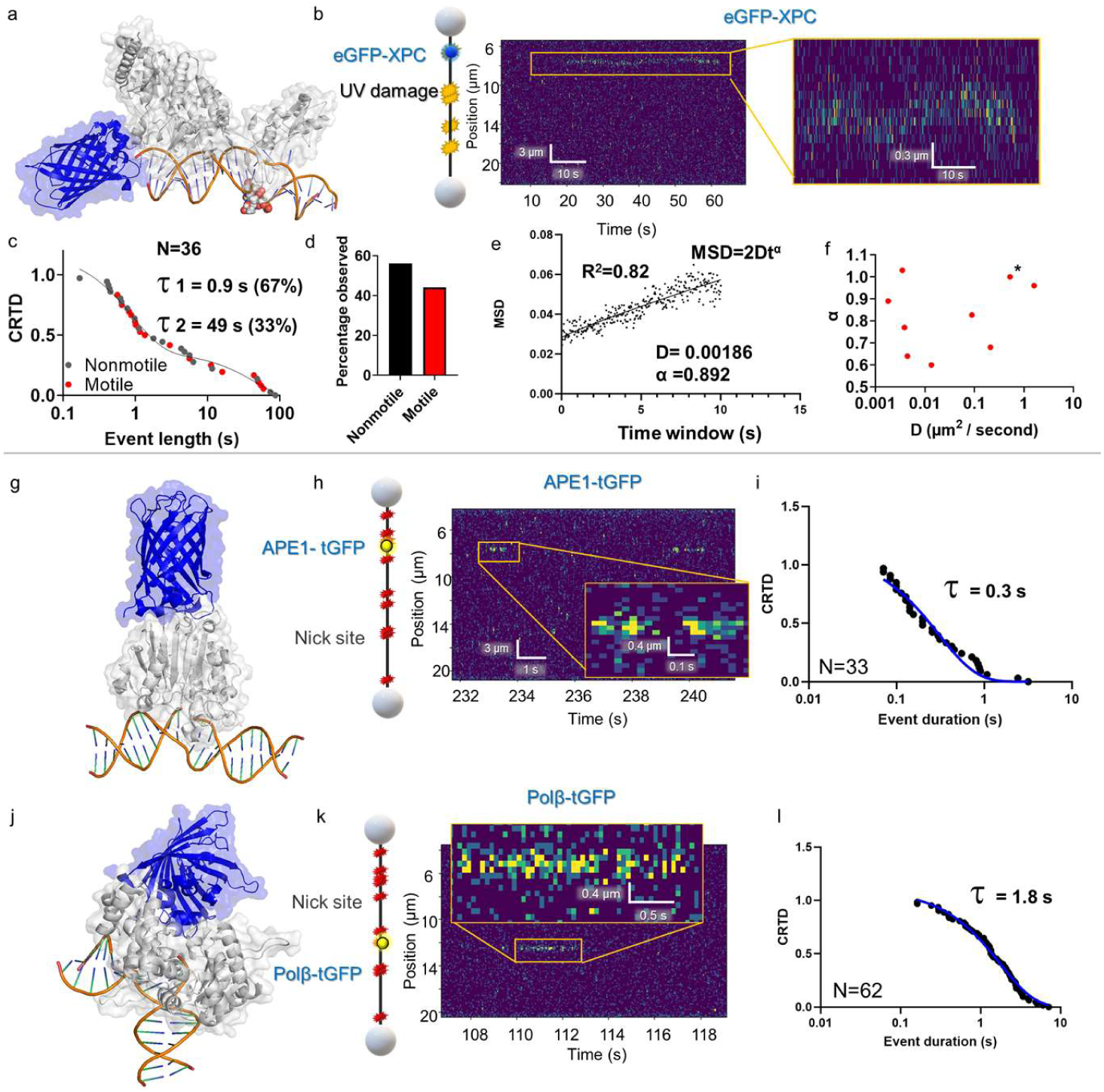
DNA-binding interactions of other DNA repair proteins. **a**, The structure of eGFP-XPC (PDB codes 6CFI of Rad4 the yeast homolog to XPC and 4EUL). **b**, a cartoon depiction of the DNA substrate used for XPC binding characterization, with UV damage sites shown in yellow and XPC binding shown in blue. Also shown is an example kymograph of eGFP-XPC binding and diffusing along the DNA in yellow. **c**, CRTD analysis of XPC binding DNA with UV damage. **d**, Distribution of motile and nomotile XPC events. **e**, An example MSD plot for analyzing XPC diffusion on DNA. **f**, Diffusion and alpha values for the diffusion of XPC on DNA. **g**, A structural model of APE1-tGFP from PDB code (5WNO and 4EUL). **h**, Schematic and example kymograph of APE1 binding to DNA with nicks. **i**, CRTD analysis of APE1 binding nicked DNA, with fit shown in blue. **j**, A structural model of pol β-tGFP, taken from PDB codes (4KLO and 4EUL) and the tGFP modeled in. **k**, Example schematics of pol β binding DNA containing nicks as well as a corresponding kymograph of an observation of pol β binding. **l**, CRTD analysis of pol β binding nicked DNA, with the fit shown in blue.

### Using SMADNE to observe protein dynamics on DNA

We next studied the DNA repair protein UV-DDB, which is composed of a heterodimer between DNA damage-binding protein 1 (DDB1, 127 kDa) and DNA damage-binding protein 2 (DDB2, 48 kDa). This latter subunit engages DNA at the site of damage^38^. UV-DDB detects UV-induced photoproducts with high affinity^39^, and the purified protein has been extensively characterized at the single-molecule level for various DNA substrates ^30,38,40^. Thus, these previous studies serve as a benchmark in which to validate the behavior of UV-DDB by SMADNE. Since UV-DDB is a heterodimer, it provided an opportunity to orthogonally label DDB1 with an N-terminal eGFP tag and DDB2 with an N-terminal HaloTag conjugated to JaneliaFluor 635 dye (Fig. 4a) ^41^.

**Fig. 4:**
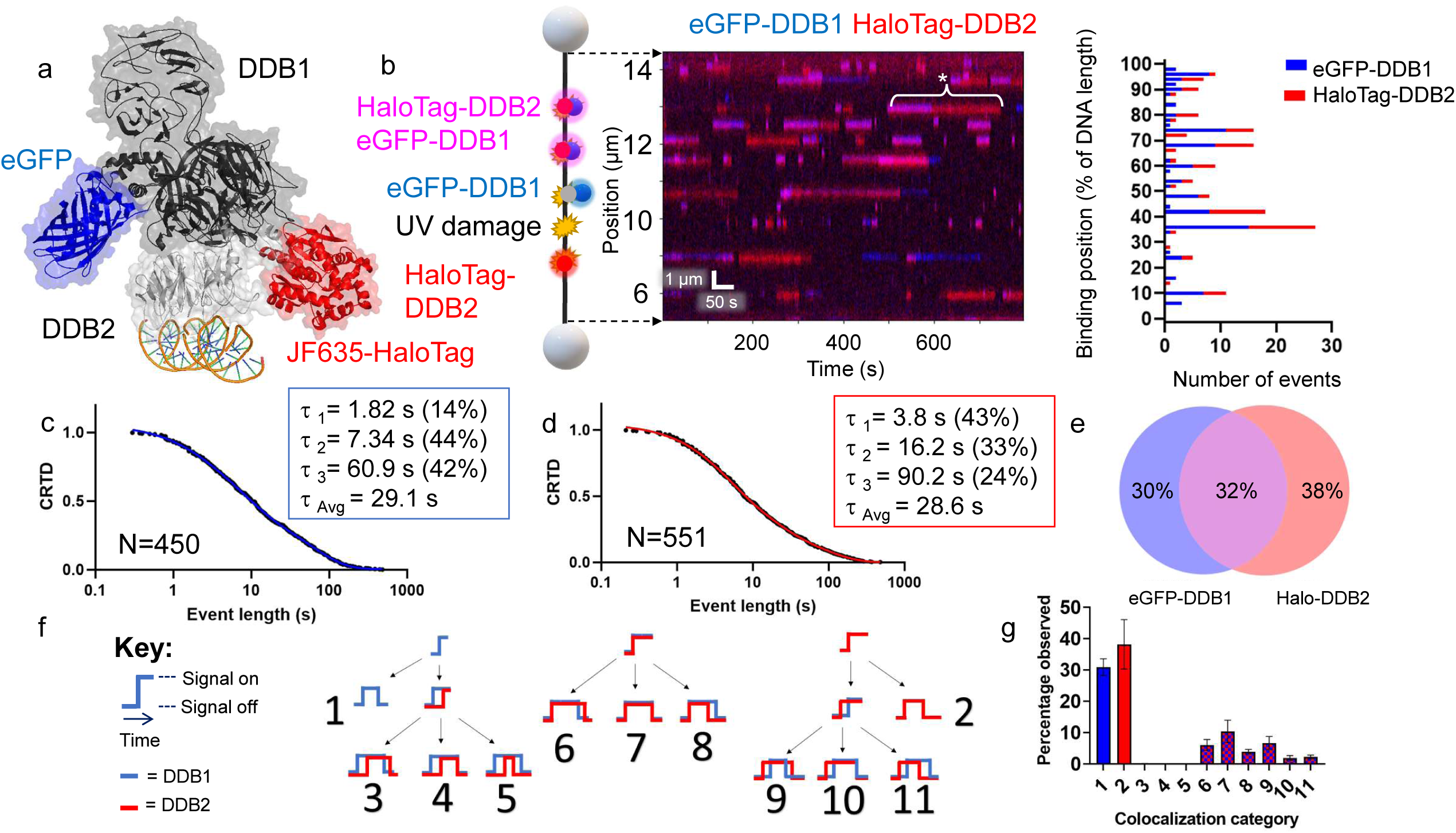
SMADNE characterization of dual-labeled UV-DDB binding UV damage. **a**, Structure of UV-DDB bound to DNA (PDB ID: 4E5Z, 4EUL, 5UY1) with modeled fluorescent tags. **b**, An example kymograph of eGFP-DDB1 (blue) and HaloTag-DDB2 (red) binding to 48.5 kb DNA with UV damage. When both colors bind together the color appears magenta. The white asterisk marks an event where DDB1 and DDB2 bound together followed by DDB1 dissociation. Also shown is a graph of the positions of events in the kymograph. **c, d** Cumulative residence time distribution (CRTD) for DDB1 (c) and DDB2 (d) binding UV-damaged DNA. **e**, Percentage of events that were DDB1 alone, DDB2 alone, or colocalized (middle). **f**, A diagram showing the 11 possible colocalization categories for two colors of molecules binding DNA. **g**, The distribution of the 11 categories for DDB1 and DDB2 binding UV-damaged DNA. Error bars represent the SEM of four experiments.

As reported by previous studies of purified protein in these buffer conditions, UV-DDB did not exhibit 1D diffusion (sliding) on the DNA but rather found its damaged substrates via 3D diffusion^38^. Furthermore, DDB1 and DDB2 bound specific positions on the DNA multiple times within a single viewing window (Fig. 4b). These long-lived binding positions (lifetimes > 10 s) presumably represent the sites of UV photoproducts after UV treatment (40 J/m^2^). Within these damage sites, some positions had many short interactions over the course of a kymograph (consistent with a low-affinity substrate being weakly bound and released multiple times) and some positions only had a few long interactions (consistent with a high-affinity substrate strongly bound by UV-DDB). This pattern may indicate binding to cyclobutane pyrimidine dimers and 6-4 photoproducts, respectively, both of which are products of UV irradiation^42^.

The binding events of both DDB1 and DDB2 exhibited a wide distribution of binding durations (four orders of magnitude) in good agreement with previous studies of purified UV-DDB (Fig. 4c, d). Binding event durations were fit to CRTD to quantify the rate of dissociation (*k*_off_) ^38^. The DDB1 and DDB2 plots were fit to a triple-exponential decay function as was previously reported for purified UV-DDB, with one short lifetime (∼2 and 4 s respectively) one medium lifetime (7 and 16 s respectively), and one long lifetime (61 and 90 s respectively) (Fig. 4c, d). The weighted average lifetime (all three lifetimes multiplied by their percentage contribution) for DDB1 was 29.1 s, relatively close to DDB2 at 28.6 s. These weighted average lifetimes were around 50% longer than the previous observations with purified UV-DDB on UV-damaged DNA (weighted average of 18.5 seconds)^38^. As the previous strategy relied on Qdot-conjugated UV-DDB, this shorter lifetime observed previously could be due to Qdot conjugation process causing a modest reduction in UV-DDB binding affinity and thus a decreased lifetime as compared to our new fusion protein approach. Alternatively, unlabeled interacting proteins in the nuclear extract, such as heat shock proteins could help stabilize UV-DDB^43^.

Using the dual-labeling approach, the frequency of DDB1 and DDB2 co-localization within the localization precision of our instrument was quantified. Consistent with UV-DDB acting as a stable heterodimer, we saw many events colocalize – 32% of events had at least one colocalization with the second color, compared to 30% of events that were either one molecule of eGFP-DDB1 or 38% that were HaloTag-DDB2 (Fig. 4e). To ensure that the colocalized binding events were from one heterodimer of UV-DDB rather than a dimer of heterodimers or two heterodimers bound closely together ^5^, we also studied a mix of two colors of HaloTag-DDB2 (JF503 and JF-635) which rarely colocalized. Since DDB2 is the subunit of UV-DDB that contains DNA-binding activity, it is possible that it could bind UV damage without DDB1. However, DDB1 does not contain a DNA-binding domain, so we hypothesize that events that appear to be DDB1 alone on the DNA are DDB1 bound to another DNA-binding protein such as an unlabeled DDB2 or other known interaction partners like CSA or CDT1, which are probably present in the extract at low concentrations ^44,45^.

Including single-color events (without colocalization), there are 11 possible event classes of molecular interactions on DNA (Fig. 4f), nine of which are colocalization events with unique assembly and disassembly mechanisms. We wrote a script to classify these 11 different types of events (publicly available on LUMICKS Harbor) and found that the most common type for our data was category 7, in which DDB1 and DDB2 arrive and dissociate together. These data are consistent with UV-DDB acting as a stable heterodimer. However, we also saw the next most common was category 9, where DDB2 binds first followed by DDB1 and then DDB1 dissociates before DDB2, suggesting that alternative modes of binding exist where the proteins sequentially assemble and disassemble from the damage. Of note, categories 3-5 appeared exceedingly rare (Fig. 4g).

### Effects of unlabeled protein on fluorescently tagged protein behavior: Facilitated dissociation of UV-DDB with purified UV-DDB

Although *k*_off_ values and thus binding lifetimes are traditionally thought to be concentration independent, a growing body of work has shown that presence of competitor proteins can alter binding lifetimes ^46-48^. This phenomenon would alter binding results observed by SMADNE if the endogenous unlabeled protein represents a significant fraction compared to the labeled protein of interest. To examine facilitated dissociation of the target labeled protein by the endogenous non-labeled protein, we included tenfold excess concentration of purified UV-DDB (3 nM) along with the eGFP-DDB1 and HaloTag-DDB2 tagged proteins in extracts (Fig. 5a). While we observed a similar number of events, the event lifetime was drastically reduced by ∼30-fold for DDB1 and ∼40-fold for DDB2 in the presence of purified protein (Fig. 5b-d). Interestingly, we also saw a decrease in colocalization frequency from 32% to 19%, which may suggest that the subunits from purified UV-DDB may exchange in solution; however, category 7 (binding together and dissociating together) was again the most common category (Fig. 5e, f).

**Fig. 5:**
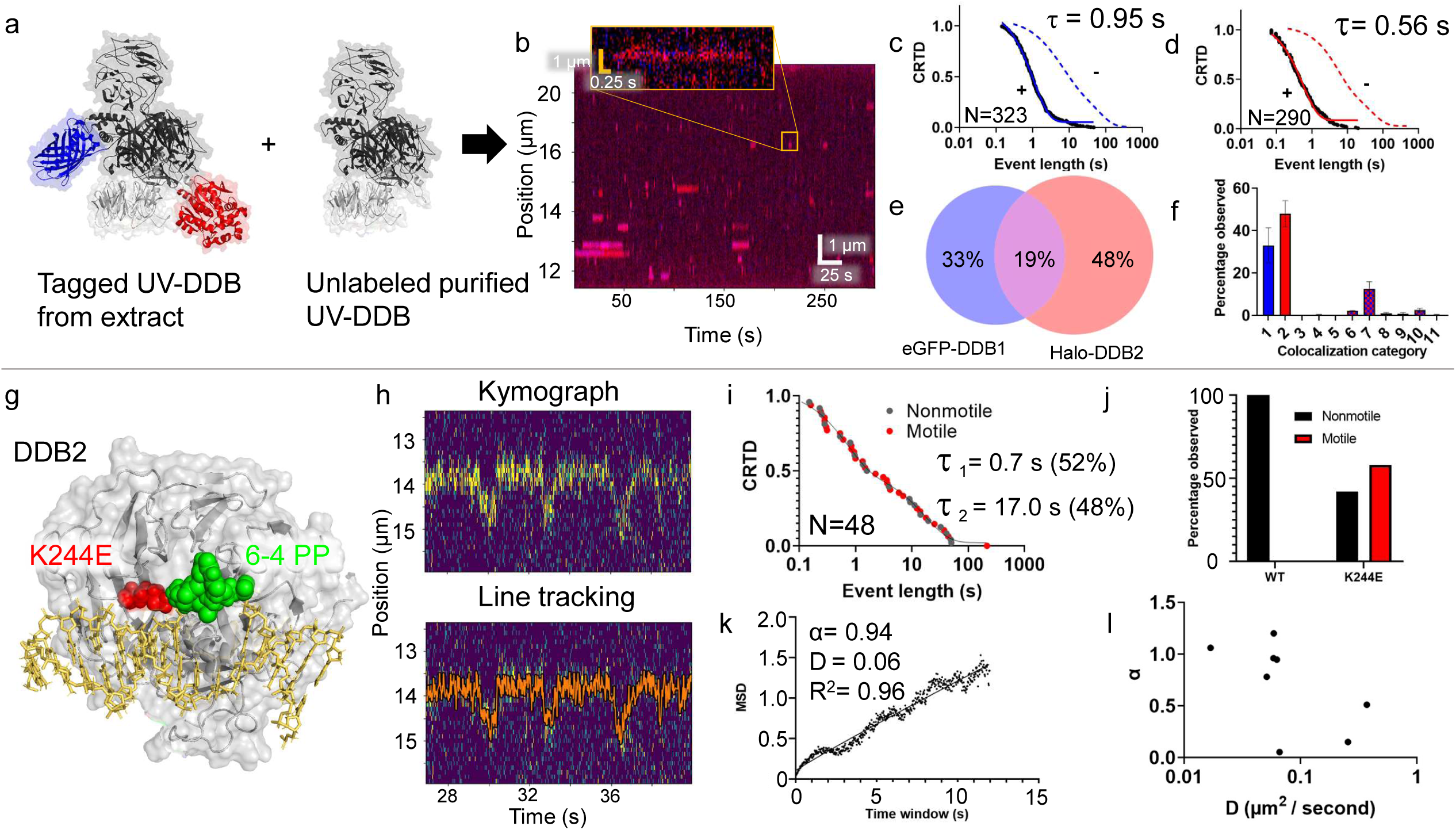
Observing facilitated dissociation and movement behavior of DDB2 K244E. **a**, Diagram of dual-labeled UV-DDB (with eGFP and HaloTag-JF-635) and unlabeled purified UV-DDB included (PDB ID: 4E5Z, 4EUL, 5UY1). **b**, An example kymograph of labeled DDB1 and DDB2 binding transiently to UV-damaged DNA. **c, d**, CRTD plots of DDB1 (blue) and DDB2 (red) shown. Dotted lines indicate CRTD curves without added unlabeled UV-DDB. **e**, Distribution of events that were DDB1 alone, DDB2 alone, or colocalized. **F**, Colocalization categories for DDB1 and DDB2 binding damaged DNA with error bars as the SEM of three experiments. **g**, Structure of DDB2 bound to a 6-4 photoproduct, with the site of the K244E mutation marked in red. **h**, A kymograph of motile DDB2 K244E binding. The tracked position of the line is shown in orange. **i**, the CRTD plot for all K244E binding events, with motile shown in red and nonmotile events shown in gray. **j**, the distribution of motile and nonmotile for WT and DDB2 K244E. **k**, Mean Squared Displacement analysis of event shown in panel h. **l**, Diffusivity (D) and α values for K244E events.

### SMADNE allows rapid characterization of protein variants: DDB2 variant (K244E)

SMADNE offers a rapid means to determine the effects of naturally occurring mutations on function without having to purify the protein, which can reduce yield and activity. To address this goal, we utilized a K244E variant of DDB2 associated with the human syndrome xeroderma pigmentosum complementation group E (Fig. 5g). Previous single-molecule characterization of this variant demonstrated this substitution causes UV-DDB to lose specificity for damage sites by diffusing past UV-induced photoproducts ^5^. Indeed, the mNeon-DDB2 K244E variant exhibited increased motility and decreased binding lifetimes (Fig. 5h), with 58% of the events observed exhibiting detectable motion in contrast to 0% with WT DDB2 (Fig. 5i, j). MSD analysis of the motile binding events indicated mNeonGreen-DDB2 K2444E behaved similarly to what we reported for Qdot labeled variant (Fig. 5k, l). One reason that the diffusivity could be slower with the purified proteins is that the Qdot label increases the drag considerably compared to the smaller fusion tag in the SMADNE approach. In addition to the motion along the DNA, shorter binding lifetimes were observed with this mutant compared to our characterization of WT DDB2, with the slowest off rate disappearing and the data was best fitted to a double exponential instead. The average lifetime for DDB2 K244E was 8.5 s, which agrees with the hypothesis that the mutation prevents full engagement with the DNA (Fig. 5l).

### Visualizing oxidative damage repair dynamics with SMADNE

Using single molecule and cellular studies we recently demonstrated that UV-DDB interacts with OGG1 to process 8-oxoG lesions^40^. To this end, we first prepared nuclear extracts from mScarlet-OGG1 expressing cells (Fig. 6a) and studied OGG1 binding on DNA treated with oxidative damage (one 8-oxoG/440 bp) ^49^. OGG1 bound to numerous positions along the length of the DNA, with many positions bound multiple times (presumably the sites of oxidative damage, Fig. 6b). Each binding position of OGG1 exhibited similar binding lifetimes: a CRTD plot revealed a best fit to a double-exponential function with a weighted average lifetime of 1.37 s (Fig. 6d). These lifetimes agree with the ∼ 2 s lifetimes published by Wallace and coworkers for purified *E*.*coli* Fpg ^49^, and Verdine and colleagues for OGG1 on non-damaged DNA^50^. We then tested the binding characteristics of a catalytically dead OGG1 variant containing a mutation in its active site, K249Q (Fig. 6c) ^51^. The binding kinetics of eGFP-labeled OGG1 K249Q on a DNA substrate containing 8-oxoG revealed much longer binding lifetimes compared to WT OGG1 (binding lifetimes of 6.2 and 36 s, with the fast lifetime contributing 75%, Fig. 6d).

**Fig. 6:**
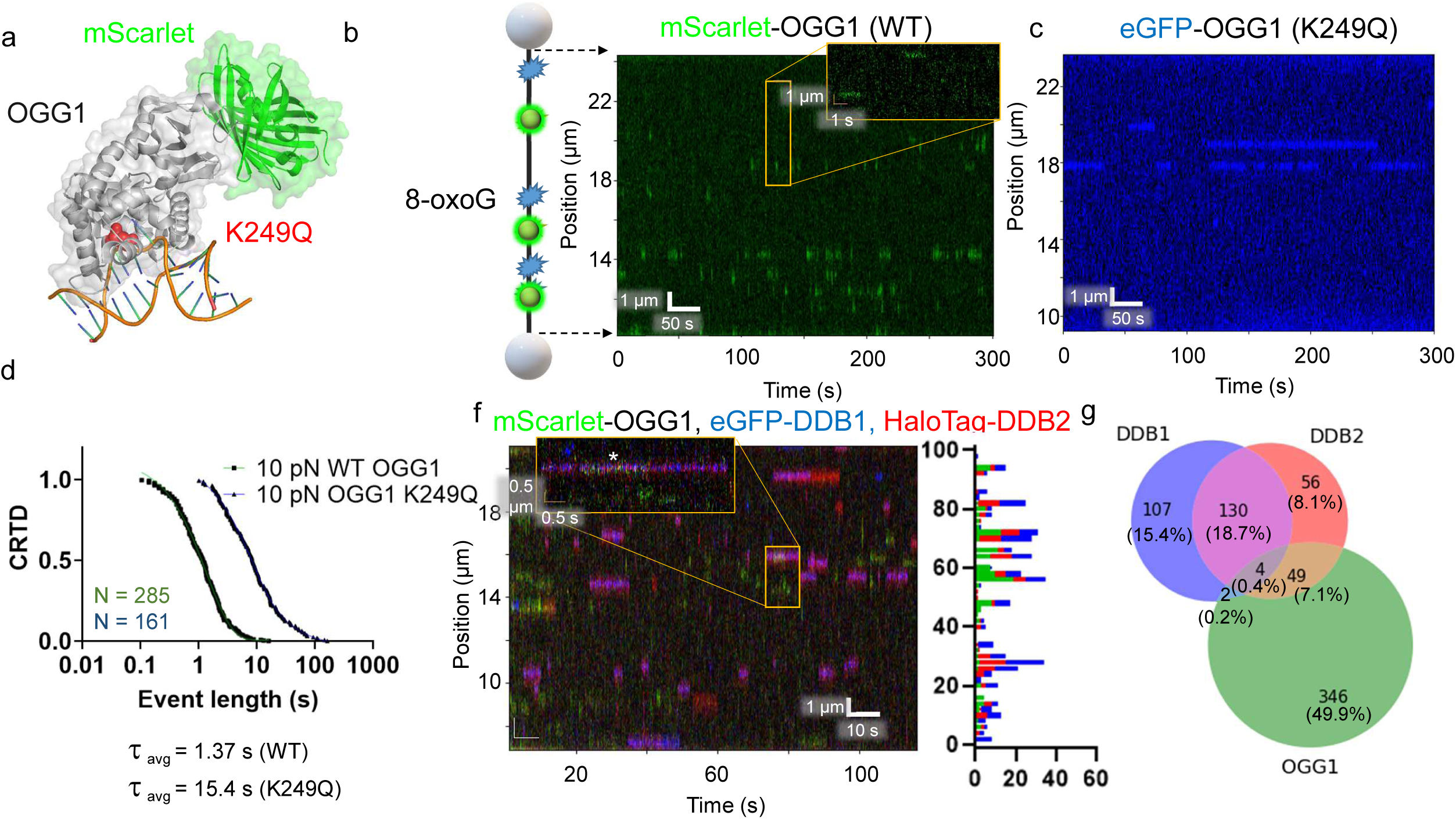
OGG1 and UV-DDB binding to DNA with oxidative damage. **a**, A structural model of mScarlet-tagged OGG1 bound to 8-oxoG containing DNA (PDB codes 1YQR and 5LK4). **b**, A cartoon depiction of DNA with 8-oxoG damage shown in blue. The accompanying kymograph it shows many transient OGG1 binding events on the DNA in green. **c**, With the catalytically dead variant K249Q, much longer binding lifetimes were observed (blue). **d**,**e**, CRTD analysis for WT and K249Q OGG1 at 1 pN and 10 pN. At 10 pN, the weighted average lifetime for the mutant was 15.4 s (42.9 and 7.7 s, 78% fast), over tenfold longer than the 1.4 s single-exponential fit. **f**, Kymograph of mScarlet-OGG1 (green), eGFP-DDB1 (blue), and HaloTag-DDB2 (red) with binding positions shown on the right. **g**, The distribution of events that bound alone vs colocalizing for all three proteins.

Since we have found that UV-DDB interacts with OGG1 to process 8-oxoG lesions^40^, we sought to determine whether these interactions could be observed in nuclear extracts using SMADNE. To this end, we expressed mScarlet-OGG1, eGFP-DDB1, and HaloTag-JF635-DDB2, and observed interactions between all three proteins (Fig. 6f, g). UV-DDB bound to DNA with oxidative damage robustly, but the binding lifetimes of DDB2 (0.14 s) were reduced compared to their lifetime on UV damage, in agreement with its lower affinity to 8-oxoG compared to UV damage^40^. Furthermore, we did see a moderate degree of transient colocalization between DDB2 and OGG1, but the majority of binding events were either OGG1 alone or DDB1 and DDB2 together at 49.9% and 15.4%, respectively (Fig. 6g).

## Discussion

SMADNE offers several major advantages compared to traditional single-molecule studies in living cells or with purified proteins. First, nuclear extracts used in SMADNE rapidly generate similar mechanistic information in agreement with previous work using purified proteins (including binding lifetimes and other outcomes shown in Fig. 1). Second, since SMADNE can utilize common fluorescence tags such as eGFP, nuclear extracts could be rapidly prepared from transfection of commercially available overexpression plasmids. Third, orthologous labeling allowed co-localization studies to be performed on heterodimeric complexes and interacting proteins. Fourth, SMADNE enables a wide range of interaction affinities to be studied, even transient interactions with *K*_D_ values of ∼1 μM. In all, the work on the UV-DDB and OGG1 variants indicate that SMADNE will provide mechanistic insights for proteins of interest via site-directed mutagenesis of specific residues.

Other methods exist that have been used to characterize proteins, RNA, and DNA at the single-molecule scale. These include Comparative Colocalization Single-Molecule Spectroscopy (CoSMoS) to study RNA-protein interactions out of yeast extracts ^18,19^, *Xenopus laevis* egg extracts to study DNA replication and repair ^20-22^ and single-molecule pulldown (SiMPull) to analyze protein complex stoichiometry and binding parameters from pulled-down proteins, among other techniques ^23-25^. SMADNE for the first time, uses human nuclear extracts to visualize protein binding on DNA strands in relation to defined genomic position and generates invaluable mechanistic information under the most physiological conditions possible. In this way post-translational modification of desired proteins after specific signaling events (e.g., DNA damage responses) can be monitored. Furthermore, performing SMADNE on the LUMICKS C-trap overcomes a disadvantage to single molecule approaches requiring TIRF microscopy that utilize DNA tethered to the bottom of the flow cells: nuclear debris can also stick to the bottom of flow chambers and obscure/overpower the fluorescence of single molecules. In contrast, with SMADNE the DNA strand remains in the center of the flow cell, circumventing debris accumulation in its focal plane. We have also found that the optical traps can additionally be used to keep the imaging zone clear from nuclear debris (see methods).

SMADNE substantially lowers the barrier of entry for numerous research groups to understand their DNA-binding proteins of interest at the single-molecule level without the burden of protein purification. While the applications shown here focus on DNA repair proteins, many other types of DNA-binding proteins would also apply, including transcription factors, helicases, and DNA polymerases. Furthermore, this new approach could be used to observe macromolecular interactions from extracts generated from a wide range of cells and tissues from animals expressing fluorescent proteins. With the rapid workflow of plasmid transfection to single-molecule data collection, SMADNE creates the possibility to screen numerous disease-associated protein variants in a high-throughput manner previously unattainable with purified proteins. Hence, SMADNE performed in conjunction with the LUMICKS C-trap represents a novel, scalable, and relatively high-throughput method to obtain single molecule mechanistic insights into key protein-DNA interactions in an environment resembling the nucleus of mammalian cells.

## Code availability

Code for converting positional data to 2D movies is available on github at https://github.com/Kad-Lab/SMADNE.

## Acknowledgements

We greatly appreciate the helpful discussions with Drs. Caroline Kisker and Jochen Kuper (University of Wurzburg), members of LUMICKS, including Aafke van den Berg and Dr. Evan Gates, as well as members of the Van Houten laboratory, including Sripriya Raja, Liam Leary, and Angela Paul. This work was supported by NIH R35ES031638 (BVH) and the Hillman Postdoctoral Fellowship for Innovative Cancer Research (MAS) and 2P30CA047904 to the UPMC Hillman Cancer Center. The manuscript’s contents are solely the responsibility of the authors and do not necessarily represent the official views of the NIEHS or NIH.

## Author contributions

B.V.H. and M.A.S. conceived the study. Nuclear extracts were prepared by N.K., V.R., R.C.J., and Z.Z. DNA substrates were generated by M.A.S. Single-molecule data was collected by M.A.S., B.L.S., R.U., and Z.Z. Analysis was performed by M.A.S., B.L.S., and R.U. with advisement from N.M.K and B.V.H. The study was written up by M.A.S. and B.V.H. with helpful input and discussion from the other authors.

## References

1 Schaich, M. A. & Van Houten, B. Searching for DNA Damage: Insights From Single Molecule Analysis. Frontiers in Molecular Biosciences 8, doi:10.3389/fmolb.2021.772877 (2021).

2 Ha, T., Kozlov, A. G. & Lohman, T. M. Single-molecule views of protein movement on single-stranded DNA. Annu Rev Biophys 41, 295–319, doi:10.1146/annurev-biophys-042910-155351 (2012).

3 Kong, M. & Greene, E. C. Mechanistic Insights From Single-Molecule Studies of Repair of Double Strand Breaks. Front Cell Dev Biol 9, 745311, doi:10.3389/fcell.2021.745311 (2021).

4 Liu, L. et al. PARP1 changes from three-dimensional DNA damage searching to one-dimensional diffusion after auto-PARylation or in the presence of APE1. Nucleic Acids Res 45, 12834–12847, doi:10.1093/nar/gkx1047 (2017).

5 Ghodke, H. et al. Single-molecule analysis reveals human UV-damaged DNA-binding protein (UV-DDB) dimerizes on DNA via multiple kinetic intermediates. Proc Natl Acad Sci U S A 111, E1862–1871, doi:10.1073/pnas.1323856111 (2014).

6 Schaich, M. A. & Van Houten, B. Searching for DNA Damage: Insights From Single Molecule Analysis. Frontiers in molecular biosciences 8, 772877, doi:10.3389/fmolb.2021.772877 (2021).

7 Kong, M., Beckwitt, E. C., Springall, L., Kad, N. M. & Van Houten, B. Single-Molecule Methods for Nucleotide Excision Repair: Building a System to Watch Repair in Real Time. Methods Enzymol 592, 213–257, doi:10.1016/bs.mie.2017.03.027 (2017).

8 Ghodke, H., Ho, H. N. & van Oijen, A. M. Single-molecule live-cell imaging visualizes parallel pathways of prokaryotic nucleotide excision repair. Nat Commun 11, 1477, doi:10.1038/s41467-020-15179-y (2020).

9 Wiktor, J. et al. RecA finds homologous DNA by reduced dimensionality search. Nature 597, 426–429, doi:10.1038/s41586-021-03877-6 (2021).

10 Wang, H., Tessmer, I., Croteau, D. L., Erie, D. A. & Van Houten, B. Functional characterization and atomic force microscopy of a DNA repair protein conjugated to a quantum dot. Nano Lett 8, 1631–1637, doi:10.1021/nl080316l (2008).

11 Gebhardt, J. C. M. et al. Single-molecule imaging of transcription factor binding to DNA in live mammalian cells. Nature Methods 10, 421–426, doi:10.1038/nmeth.2411 (2013).

12 Liu, H. et al. Visualizing long-term single-molecule dynamics in vivo by stochastic protein labeling. Proceedings of the National Academy of Sciences 115, 343, doi:10.1073/pnas.1713895115 (2018).

13 Betzig, E. et al. Imaging intracellular fluorescent proteins at nanometer resolution. Science 313, 1642–1645, doi:10.1126/science.1127344 (2006).

14 Hess, S. T., Girirajan, T. P. K. & Mason, M. D. Ultra-high resolution imaging by fluorescence photoactivation localization microscopy. Biophys J 91, 4258–4272, doi:10.1529/biophysj.106.091116 (2006).

15 Rust, M. J., Bates, M. & Zhuang, X. Sub-diffraction-limit imaging by stochastic optical reconstruction microscopy (STORM). Nat Methods 3, 793–795, doi:10.1038/nmeth929 (2006).

16 Elf, J., Li, G.-W. & Xie, X. S. Probing Transcription Factor Dynamics at the Single-Molecule Level in a Living Cell. Science 316, 1191–1194, doi:10.1126/science.1141967 (2007).

17 Schmidt, J. C., Zaug, A. J. & Cech, T. R. Live Cell Imaging Reveals the Dynamics of Telomerase Recruitment to Telomeres. Cell 166, 1188-1197.e1189, doi:10.1016/j.cell.2016.07.033 (2016).

18 Haraszti, R. A. & Braun, J. E. Comparative Colocalization Single-Molecule Spectroscopy (CoSMoS) with Multiple RNA Species. Methods Mol Biol 2113, 23–29, doi:10.1007/978-1-0716-0278-2_3 (2020).

19 Hoskins, A. A. et al. Ordered and dynamic assembly of single spliceosomes. Science (New York, N.Y.) 331, 1289–1295, doi:10.1126/science.1198830 (2011).

20 Sparks, J. L. et al. The CMG Helicase Bypasses DNA-Protein Cross-Links to Facilitate Their Repair. Cell 176, 167-181.e121, doi:10.1016/j.cell.2018.10.053 (2019).

21 Kanke, M., Tahara, E., Huis In’t Veld, P. J. & Nishiyama, T. Cohesin acetylation and Wapl-Pds5 oppositely regulate translocation of cohesin along DNA. Embo j 35, 2686–2698, doi:10.15252/embj.201695756 (2016).

22 Graham, T. G. W., Walter, J. C. & Loparo, J. J. Two-Stage Synapsis of DNA Ends during Non-homologous End Joining. Mol Cell 61, 850–858, doi:10.1016/j.molcel.2016.02.010 (2016).

23 Aggarwal, V. & Ha, T. Single-molecule pull-down (SiMPull) for new-age biochemistry. BioEssays 36, 1109–1119, doi:https://doi.org/10.1002/bies.201400090 (2014).

24 Jain, A., Liu, R., Xiang, Y. K. & Ha, T. Single-molecule pull-down for studying protein interactions. Nat Protoc 7, 445–452, doi:10.1038/nprot.2011.452 (2012).

25 Jain, A. et al. Probing cellular protein complexes using single-molecule pull-down. Nature 473, 484–488, doi:10.1038/nature10016 (2011).

26 Lee, J.-B. et al. DNA primase acts as a molecular brake in DNA replication. Nature 439, 621–624, doi:10.1038/nature04317 (2006).

27 Roy, R., Kozlov, A. G., Lohman, T. M. & Ha, T. SSB protein diffusion on single-stranded DNA stimulates RecA filament formation. Nature 461, 1092–1097, doi:10.1038/nature08442 (2009).

28 Gorman, J. & Greene, E. C. Visualizing one-dimensional diffusion of proteins along DNA. Nature Structural & Molecular Biology 15, 768–774, doi:10.1038/nsmb.1441 (2008).

29 Qi, Z. et al. DNA Sequence Alignment by Microhomology Sampling during Homologous Recombination. Cell 160, 856–869, doi:https://doi.org/10.1016/j.cell.2015.01.029 (2015).

30 Jang, S. et al. Single molecule analysis indicates stimulation of MUTYH by UV-DDB through enzyme turnover. Nucleic Acids Res 49, 8177–8188, doi:10.1093/nar/gkab591 (2021).

31 Hughes, C. D. et al. Real-time single-molecule imaging reveals a direct interaction between UvrC and UvrB on DNA tightropes. Nucleic Acids Research 41, 4901–4912, doi:10.1093/nar/gkt177 (2013).

32 Rad, B., Forget, A. L., Baskin, R. J. & Kowalczykowski, S. C. Single-molecule visualization of RecQ helicase reveals DNA melting, nucleation, and assembly are required for processive DNA unwinding. Proceedings of the National Academy of Sciences 112, E6852, doi:10.1073/pnas.1518028112 (2015).

33 Graham, J. E., Marians, K. J. & Kowalczykowski, S. C. Independent and Stochastic Action of DNA Polymerases in the Replisome. Cell 169, 1201-1213.e1217, doi:10.1016/j.cell.2017.05.041 (2017).

34 Choi, P. J., Cai, L., Frieda, K. & Xie, X. S. A stochastic single-molecule event triggers phenotype switching of a bacterial cell. Science (New York, N.Y.) 322, 442–446, doi:10.1126/science.1161427 (2008).

35 Liu, T.-C. et al. APE1 distinguishes DNA substrates in exonucleolytic cleavage by induced space-filling. Nature Communications 12, 601, doi:10.1038/s41467-020-20853-2 (2021).

36 Cheon, N. Y., Kim, H.-S., Yeo, J.-E., Schärer, O. D. & Lee, J. Y. Single-molecule visualization reveals the damage search mechanism for the human NER protein XPC-RAD23B. Nucleic Acids Research 47, 8337–8347, doi:10.1093/nar/gkz629 (2019).

37 Freudenthal, B. D., Beard, W. A., Shock, D. D. & Wilson, S. H. Observing a DNA polymerase choose right from wrong. Cell 154, 157–168, doi:10.1016/j.cell.2013.05.048 (2013).

38 Ghodke, H. et al. Single-molecule analysis reveals human UV-damaged DNA-binding protein (UV-DDB) dimerizes on DNA via multiple kinetic intermediates. Proceedings of the National Academy of Sciences 111, E1862, doi:10.1073/pnas.1323856111 (2014).

39 Fujiwara, Y. et al. Characterization of DNA recognition by the human UV-damaged DNA-binding protein. J Biol Chem 274, 20027–20033, doi:10.1074/jbc.274.28.20027 (1999).

40 Jang, S. et al. Damage sensor role of UV-DDB during base excision repair. Nat Struct Mol Biol 26, 695–703, doi:10.1038/s41594-019-0261-7 (2019).

41 Los, G. V. et al. HaloTag: a novel protein labeling technology for cell imaging and protein analysis. ACS Chem Biol 3, 373–382, doi:10.1021/cb800025k (2008).

42 Lo, H.-L. et al. Differential biologic effects of CPD and 6-4PP UV-induced DNA damage on the induction of apoptosis and cell-cycle arrest. BMC Cancer 5, 135–135, doi:10.1186/1471-2407-5-135 (2005).

43 Zou, Y., Crowley, D. J. & Van Houten, B. Involvement of Molecular Chaperonins in Nucleotide Excision Repair: DnaK LEADS TO INCREASED THERMAL STABILITY OF UvrA, CATALYTIC UvrB LOADING, ENHANCED REPAIR, AND INCREASED UV RESISTANCE*. Journal of Biological Chemistry 273, 12887–12892, doi:https://doi.org/10.1074/jbc.273.21.12887 (1998).

44 Fischer, E. S. et al. The molecular basis of CRL4DDB2/CSA ubiquitin ligase architecture, targeting, and activation. Cell 147, 1024–1039, doi:10.1016/j.cell.2011.10.035 (2011).

45 Cho Nathan, H. et al. OpenCell: Endogenous tagging for the cartography of human cellular organization. Science 375, eabi6983, doi:10.1126/science.abi6983.

46 Graham, J. S., Johnson, R. C. & Marko, J. F. Concentration-dependent exchange accelerates turnover of proteins bound to double-stranded DNA. Nucleic Acids Res 39, 2249–2259, doi:10.1093/nar/gkq1140 (2011).

47 Ha, T. Single-molecule approaches embrace molecular cohorts. Cell 154, 723–726, doi:10.1016/j.cell.2013.07.012 (2013).

48 Gibb, B. et al. Concentration-dependent exchange of replication protein A on single-stranded DNA revealed by single-molecule imaging. PLoS One 9, e87922, doi:10.1371/journal.pone.0087922 (2014).

49 Nelson, S. R., Dunn, A. R., Kathe, S. D., Warshaw, D. M. & Wallace, S. S. Two glycosylase families diffusively scan DNA using a wedge residue to probe for and identify oxidatively damaged bases. Proceedings of the National Academy of Sciences 111, E2091, doi:10.1073/pnas.1400386111 (2014).

50 Blainey, P. C., van Oijen, A. M., Banerjee, A., Verdine, G. L. & Xie, X. S. A base-excision DNA-repair protein finds intrahelical lesion bases by fast sliding in contact with DNA. Proceedings of the National Academy of Sciences 103, 5752, doi:10.1073/pnas.0509723103 (2006).

51 Nash, H. M., Lu, R., Lane, W. S. & Verdine, G. L. The critical active-site amine of the human 8-oxoguanine DNA glycosylase, hOgg1: direct identification, ablation and chemical reconstitution. Chemistry & biology 4, 693–702, doi:10.1016/s1074-5521(97)90225-8 (1997).

